# Progenitor derived glia are required for spinal cord regeneration in zebrafish

**DOI:** 10.1101/2022.08.24.505140

**Authors:** Lili Zhou, Ryan McAdow, Hunter Yamada, Brooke Burris, Dana Klatt Shaw, Kelsey Oonk, Kenneth D. Poss, Mayssa H. Mokalled

**Affiliations:** Department of Developmental Biology, Washington University School of Medicine, St. Louis, MO, USA; Center of Regenerative Medicine, Washington University School of Medicine, St. Louis, MO, USA; Duke Regeneration Center, Duke University Medical Center, Durham, NC, USA

## Abstract

Unlike mammals, adult zebrafish undergo spontaneous recovery after major spinal cord injury. Whereas reactive gliosis presents a roadblock for mammalian spinal cord repair, glial cells in zebrafish elicit pro-regenerative bridging functions after injury. Here, we perform genetic lineage tracing, assessment of regulatory sequences, and inducible cell ablation to define mechanisms that direct the molecular and cellular responses of glial cells after spinal cord injury in adult zebrafish. Using a newly generated CreER^T2^ transgenic line, we show that cells that direct expression of the bridging glial marker *ctgfa* give rise to regenerating glia after injury, with negligible contribution to either neuronal or oligodendrocyte lineages. A 1 kb sequence upstream of the *ctgfa* gene was sufficient to direct expression in early bridging glia after injury. Finally, ablation of *ctgfa*-expressing cells using a transgenic nitroreductase strategy impaired glial bridging and recovery of swim behavior after injury. This study identifies key regulatory features, cellular progeny, and requirements of glial cells during innate spinal cord regeneration.

## INTRODUCTION

Disparate glial, neuronal, and systemic injury responses underlie differential regenerative capacities of CNS tissues across vertebrates (Silver and Miller, 2004; Mokalled et al., 2016; Silver, 2016; O’Shea et al., 2017; Brennan and Popovich, 2018; Sofroniew, 2018). While spinal cord injuries (SCI) cause irreversible sensory and motor deficits in mammals (He and Jin, 2016; Silver, 2016; Sofroniew, 2018), adult zebrafish possess potent regenerative capacity and reverse paralysis within 8 weeks of complete spinal cord (SC) transection (Becker et al., 1997; Reimer et al., 2008; Goldshmit et al., 2012). Following SCI, mammalian astrocytes display multifarious injury responses that are overshadowed by anti-regenerative scar-forming cells and inhibitory extracellular molecules (Grimpe and Silver, 2004; Dias and Goritz, 2018; Sofroniew, 2018). Though the complexities of glial cell responses to injury have been extensively studied in mammals, our understanding of how glial cells respond to SCI in highly regenerative vertebrates like zebrafish is relatively limited. Following SCI in zebrafish, specialized glial cells form a bridge that is thought to provide a physical and signaling scaffold for cellular regrowth across the lesion (Goldshmit et al., 2012; Mokalled et al., 2016). However, whether glial cells are pro-regenerative in zebrafish and how pro-regenerative glia support innate SC regeneration require further investigation.

Previous studies identified *connective tissue growth factor a* (*ctgfa*) as a central glial bridging factor and described the emergence of *ctgfa*^*+*^ *gfap*^*+*^ bridging glia after SCI (Mokalled et al., 2016). Ctgf is a multidomain-containing extracellular molecule that interacts with multiple signaling pathways and elicits a range of cellular responses including cell proliferation and differentiation (Grotendorst and Duncan, 2005). The spatiotemporal pattern of *ctgfa* expression correlates with glial bridge formation in zebrafish (Mokalled et al., 2016). Following SCI, *ctgfa* expression is first broadly induced in proliferating *sox2*^*+*^ progenitors around the central canal proximal to the lesion. During subsequent steps of regeneration, *ctgfa* transcripts localize to a subset of ventral progenitors that undergo epithelial-to-mesenchymal transition (EMT) after injury (Mokalled et al., 2016; Klatt Shaw and Mokalled, 2021; Klatt Shaw et al., 2021). Genetic evidence indicates that *ctgfa* functions within a multi-nodal EMT-driving gene regulatory network that is necessary and sufficient to reprogram pro-regenerative glia after injury (Klatt Shaw et al., 2021). Here, we employ a suite of *ctgf*-based genetic tools to trace the fates and functions of pro-regenerative glia during SC regeneration.

Zebrafish glia possess an astrocyte-like cell identity. However, EMT gene expression distinguishes zebrafish bridging glia from mammalian astrocytes (Klatt Shaw et al., 2021). EMT is often linked to increased plasticity and stem cell activation during tissue regeneration (Jessen and Arthur-Farraj, 2019; Wilson et al., 2020), suggesting zebrafish glia have increased EMT-mediated plasticity and regenerative potential. In the zebrafish SC, EMT activation during bridging involves Yap/Taz, Egr1, Junbb, and Spi1 transcriptional regulators (Klatt Shaw et al., 2021). Hallmarks of EMT include downregulation of epithelial markers like E-cadherin and upregulation of mesenchymal genes like N-cadherin, driven by Twist and Zeb transcription factors (Dongre and Weinberg, 2019). Injury-induced expression of *egr1* and *junbb*, along with *yap/taz* activation, correlate with *ctgfa* expression and are required for bridging and functional SC repair (Gervasi et al., 2012; Wu et al., 2017; Han et al., 2019; Sato et al., 2020). However, the temporal and transcriptional hierarchies of EMT driving transcriptional regulators and the mechanisms that direct glial progenitors toward specific cell fates during regeneration remain to be dissected.

This study defines glial cell functions during zebrafish SC regeneration. Using a battery of *ctgfa*-based genetic tools, we determined the contribution, regulation and requirement of bridging glial cells during innate SC repair. A newly generated *ctgfa:*CreER^T2^ transgenic line established the contribution of *ctgfa* expressing cells to regenerative glia and glial progenitors after injury. Using a series of transgenic reporter lines, we found that regulation of *ctgfa* expression converges onto an EMT-driving distal enhancer element that directs *ctgfa-*dependent transcription following injury. Finally, we show that genetic ablation of *ctgfa*-expressing cells impairs glial bridging and functional SC repair. Our findings indicate the cell contributions and regulatory mechanisms that direct regenerative gliogenesis during innate SC repair.

## RESULTS

### Generation of *ctgfa:*CreER^T2^ zebrafish for genetic lineage tracing

Using *-5*.*5Kb-ctgfa:*EGFP and *-5*.*5Kb-ctgfa:*mCherry-NTR reporter lines, we previously showed that *ctgfa*-driven EGFP fluorescence recapitulates endogenous mRNA expression in ventral ependymal progenitors and in bridging glia during SC regeneration (Mokalled et al., 2016; Klatt Shaw et al., 2021). To specifically target *ctgfa*^*+*^ cells for permanent labeling and lineage tracing, we generated a *-5*.*5Kb-ctgfa:*CreER^T2^ line to use in combination with a previously established *ubiquitin:*loxP-GFP-STOP-loxP-mCherry transgene (Mosimann et al., 2011). The compound transgenic line *-5*.*5Kb-ctgfa:*CreER^T2^;*ubiquitin:*loxP-GFP-STOP-loxP-mCherry, referred to hereafter as *ctgfa*-Tracer, enables permanent mCherry labeling in *ctgfa*^+^ cells and their progeny following Tamoxifen-induced Cre-mediated recombination (Fig. S1A). To this end, we performed SC transections on *ctgfa*-Tracer animals and control siblings, and treated with 5 mM Tamoxifen at 4 dpi for 24 hrs to induce recombination prior to the emergence of bridging glia. To assess the extent of recombination, we collected SC tissues 2 days following the end of Tamoxifen treatment, which corresponds to 7 dpi. By EGFP (*ctgfa*-Tracer^-^) and mCherry (*ctgfa*-Tracer^+^) staining, 5% of the cells within SC tissues showed recombined *ctgfa*-Tracer^+^ expression in Tamoxifen-treated CreER^T2+^ animals (Fig. S1B,C). Conversely, SC sections from vehicle-treated CreER^T2+^, vehicle-treated CreER^T2-^, or Tamoxifen-treated CreER^T2-^ controls did not show recombined *ctgfa*-Tracer^+^ cells (Fig. S1B,C). These studies showed that recombination in the *ctgfa*-Tracer line is both Cre-dependent and Tamoxifen-inducible, and established a recombination paradigm that recapitulates the expression of endogenous *ctgfa* transcripts and of previously established *ctgfa* reporter lines after SCI (Mokalled et al., 2016; Klatt Shaw et al., 2021). We thus used the *ctgfa*-Tracer line to map the contributions of *ctgfa*^+^ cells during SC regeneration.

### Contribution of *ctgfa*^+^ cells to regenerating glia during SC regeneration

To examine the fates of *ctgfa*^+^ cells during SC regeneration, we performed SCI on *ctgfa*-Tracer animals, treated with Tamoxifen at 4 dpi and traced recombined cells at 7, 14 and 28 dpi (Fig. 1A). SC sections 150 µm (proximal) and 450 µm (distal) rostral to the lesion were analyzed using mCherry staining to label recombined *ctgfa*-Tracer^+^ cells, and Gfap staining for glial cell co-labeling (Fig. 1B). At 7 dpi, we observed infrequent Gfap^+^ projections that were *ctgfa*-Tracer^+^ proximal to the lesion, and weak Gfap^+^ *ctgfa*-Tracer^+^ signal ventral to the central canal in distal SC sections. At 14 dpi, Gfap^+^ *ctgfa*-Tracer^+^ cells were more abundant in SC tissues proximal to the lesion, and showed consistent labeling in progenitor cells ventral to the central canal in distal SC sections. By 28 dpi, the majority of *ctgfa*-Tracer^+^ cells within SC tissues were Gfap^+^ and localized to the ventral progenitors at the proximal and distal levels. We noted that immunostaining signals for filamentous Gfap intermediate filaments do not allow for exact counts of glial cell soma, and result in underestimated assessment of co-localization. Despite these limitations, quantification of Gfap^+^ *ctgfa*-Tracer^+^ co-localization indicated increased Gfap expression in recombined cells at 14 and 28 dpi relative to 7 dpi at 150 and 450 µm rostral to the lesion (Fig. 1C). For instance, at 150 µm rostral to the lesion, Gfap expression increased from 7.7% of *ctgfa*-Tracer^+^ cells at 7 dpi to 25.5% of *ctgfa*-Tracer^+^ cells at 28 dpi (Fig. 1C). Conversely, the proportions of Gfap^+^ *ctgfa*-Tracer^+^ co-localization relative to Gfap^+^ did not significantly change between 7 and 28 dpi (Fig. 1D), ostensibly due to increased Gfap expression in proximal tissues as SC tissues regenerate. Gfap^+^ *ctgfa*-Tracer^+^ fluorescence at 750 µm rostral to the lesion was unchanged across time points, suggesting the distal contribution of *ctgfa*-Tracer cells is minimal (Fig. 1C,D). These experiments indicated increased contribution of *ctgfa*-derived cells to bridging glia and ventral ependymal progenitors during SC regeneration.

**Figure 1.**
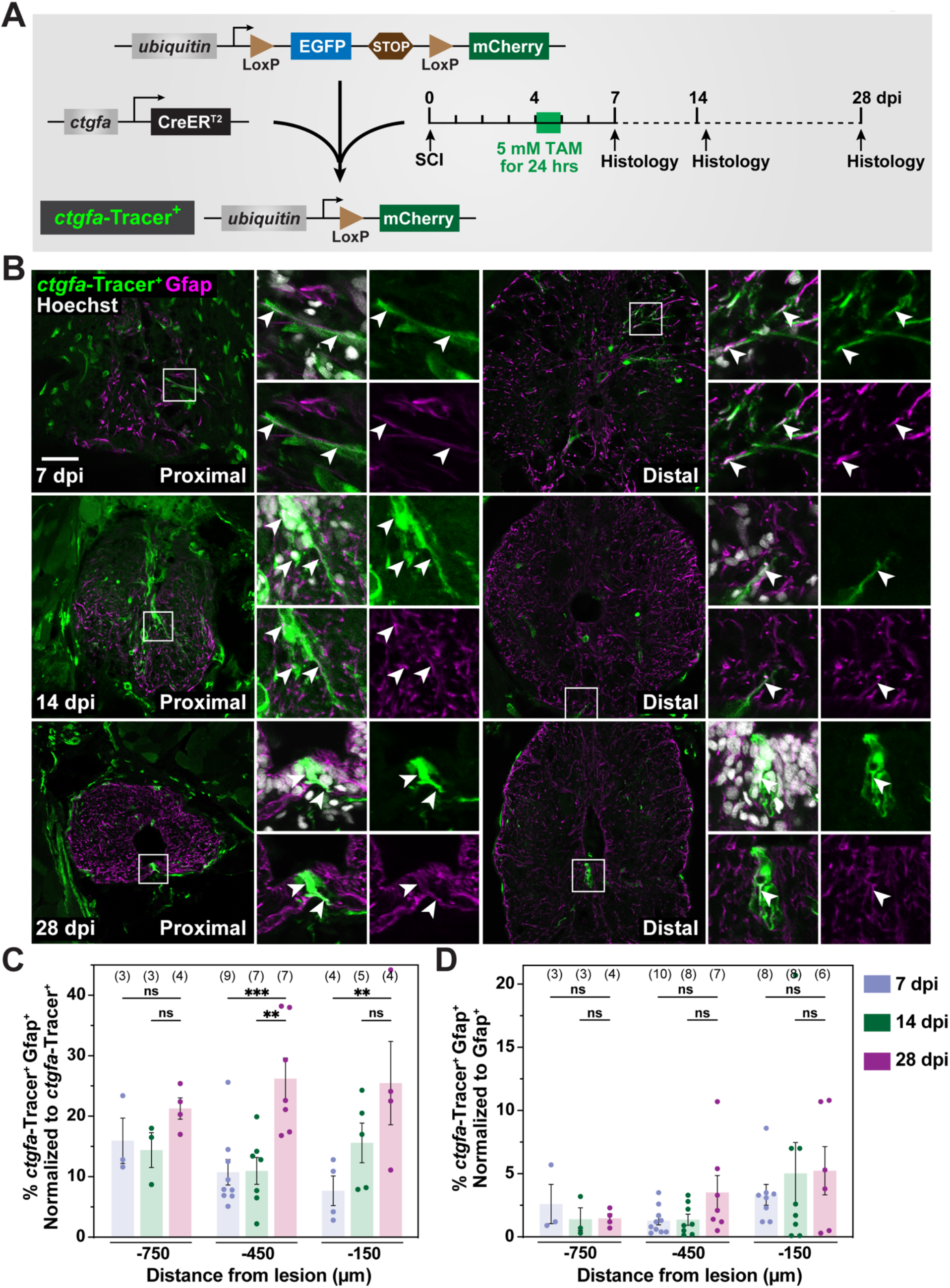
Contribution of *ctgfa*^+^ cells to regenerating glia during SC repair. **(A)** Experimental timeline to evaluate the contribution of *ctgfa*^+^ cells after SCI. *ctgfa*-Tracer animals refer to the compound transgenic line *-5*.*5Kb-ctgfa:*CreER^T2^; *ubiquitin:*loxP-GFP-STOP-loxP-mCherry. *ctgfa*-Tracer fish were subjected to complete SC transection and Tamoxifen (TAM) mediated recombination at 4 dpi to enable permanent mCherry labeling in *ctgfa*^+^ and *ctgfa*^+^-derived cells. SC tissues were collected at 7, 14 and 28 dpi for histological examination. The 7 dpi time point was used to assess recombination following TAM treatment. The 14 and 28 dpi time points were used to trace the fates of *ctgfa*^+^-derived cells. **(B)** Immunostaining for mCherry (green), Gfap (magenta), and Hoechst (grey) at 7, 14, and 28 dpi. mCherry expression, referred to as *ctgfa*-Tracer^+^, is used to trace the fates of *ctgfa* expressing cells following TAM inducible recombination. SC sections from TAM-treated *ctgfa*-Tracer (Cre^+^) animals are shown. Cross sections 150 µm (Proximal) or 450 µm (Distal) rostral from the lesion site are shown. High magnification insets show select *ctgfa*-Tracer^+^ cells in triple, double, or single channel views. Arrowheads indicate *ctgfa*-Tracer^+^ cells. **(C**,**D)** Quantification of *ctgfa*-Tracer^+^ and Gfap^+^ colocalization at 7, 14, and 28 dpi. SC cross sections 150, 450, and 750 µm rostral to the lesion were analyzed. *ctgfa*-Tracer^+^ and Gfap^+^ fluorescence were quantified. For each section, percent *ctgfa*-Tracer^+^ Gfap^+^ fluorescence was normalized to total *ctgfa*-Tracer^+^ in C and to total Gfap^+^ in D. Dots indicate individual animals and sample sizes are indicated in parentheses. ***P<0.001; **P<0.01; ns indicates P>0.05. Scale bars, 50 µm.

### Contribution of *ctgfa*^+^ cells to neurons and oligodendrocytes during SC regeneration

To evaluate the contribution of *ctgfa*^*+*^ cells to additional regenerative cell fates after SCI, we traced and quantified *ctgfa*-derived neurons and motor neurons at 28 dpi. Using the same recombination paradigm described above, we performed mCherry staining to label recombined *ctgfa*-Tracer^+^ cells and co-stained with either HuC/D to label post-mitotic neurons (Fig. 2A) or Hb9 to label newly formed motor neurons (Fig. S2A). By immunostaining, nuclear and cytoplasmic mCherry signals mark *ctgfa*-Tracer^+^ cells. Similarly, HuC/D antigens are detected in the cytoplasms and nuclei of postmitotic neurons, whereas Hb9 is eclusively nuclear in newly formed motor neurons. This allowed us to quantify the numbers of *ctgfa*-Tracer^+^ HuC/D^+^ Hoechst^+^ cells and *ctgfa*-Tracer^+^ Hb9^+^ Hoechst^+^ cells for these experiments. At 28 dpi, we occasionally observed up to 3 HuC/D^+^ *ctgfa*-Tracer^+^ cells per section (Fig. 2B, S2B) or a single Hb9^+^ *ctgfa*-Tracer^+^ cell per section (Fig. S2C,D), suggesting the contribution of *ctgfa*-Tracer cells to regenerating neurons or motor neurons were negligible. At 150 µm rostral to the lesion, the HuC/D^+^ *ctgfa*-Tracer^+^ cells accounted for 4% of *ctgfa*-Tracer^+^ cells and 5% of HuC/D^+^ neurons (Fig. 2C,D). On the other hand, Hb9^+^ neurons were not detected in any sections at either 150 or 450 µm rostral to the lesion at 28 dpi (Fig. S2E,F). We next assessed the contribution of *ctgfa*^*+*^ cells to myelinating oligodendrocytes by co-labeling of *ctgfa*-Tracer cells with Myelin basic proteins (Mbp) (Fig. 2E). Like Gfap, immunostaining for Mbp does not allow for exact counts of oligodendrocyte cells. We thus quantified Mbp^+^ *ctgfa*-Tracer^+^ co-localization, which accounted for 10.8% of *ctgfa*-Tracer^+^ (Fig. 2F), and for <0.3% of Mbp^+^ fluorescence (Fig. 2G). These studies indicated the contribution of *ctgfa*^+^ cells to regenerating glial cells, with minimal contribution to neuron or oligodendrocyte lineages during SC regeneration.

**Figure 2.**
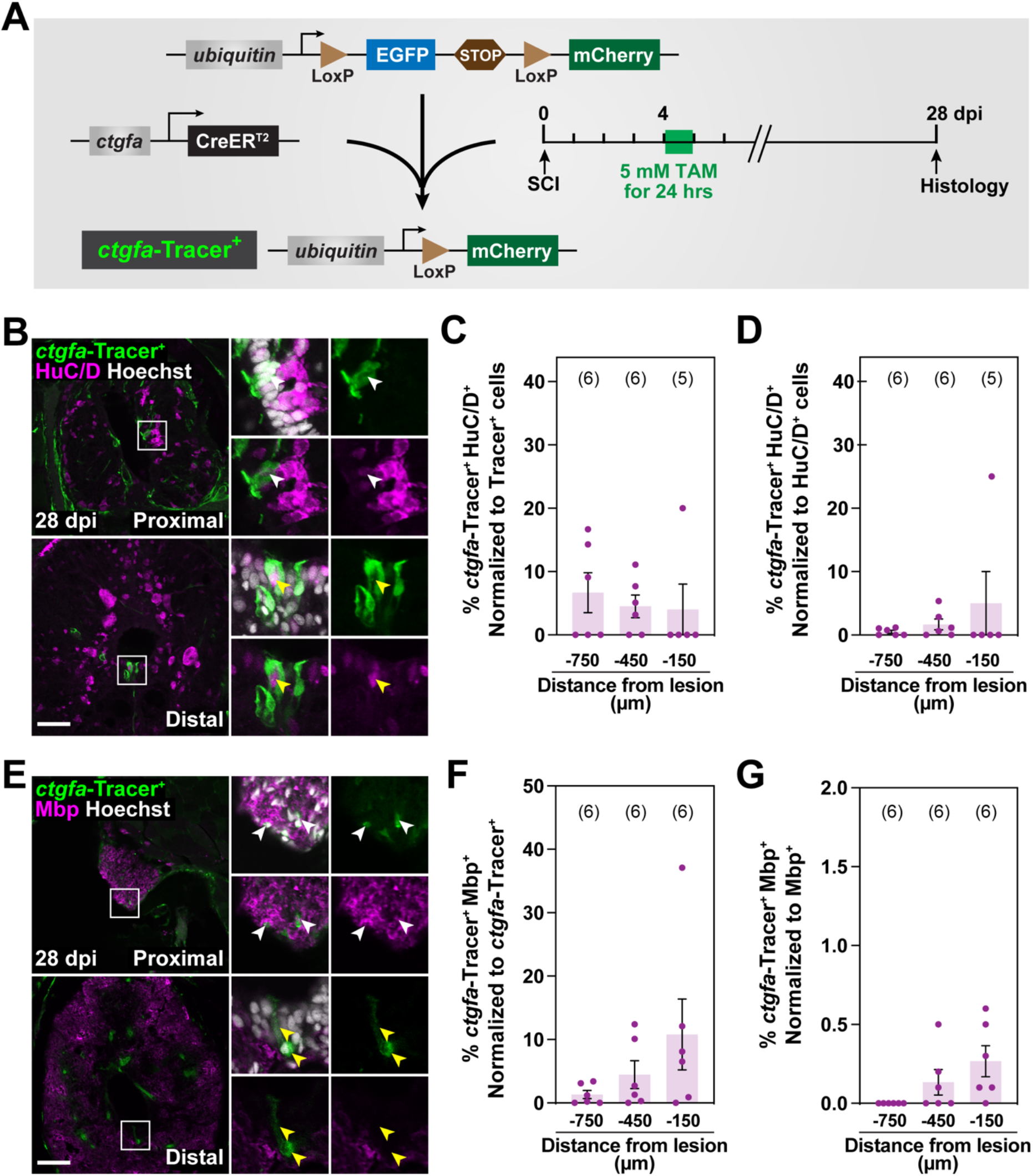
Contribution of *ctgfa*^+^ cells to regenerating neurons and oligodendrocytes after SCI. **(A)** Experimental timeline to evaluate the contribution of *ctgfa*^+^ cells after SCI. *ctgfa*-Tracer animals were subjected to complete SC transection and Tamoxifen (TAM) mediated recombination at 4 dpi to enable permanent mCherry labeling in *ctgfa*^+^ and *ctgfa*^+^-derived cells. SC tissues were collected at 28 dpi for histological examination. **(B)** Immunostaining for mCherry (green), HuC/D (magenta), and Hoechst (grey) at 28 dpi. mCherry expression, referred to as *ctgfa*-Tracer^+^, is used to trace the fates of *ctgfa* expressing cells following TAM inducible recombination at 4 dpi. SC sections from TAM-treated *ctgfa*-Tracer (Cre^+^) animals are shown. Cross sections 150 µm (Proximal) or 450 µm (Distal) rostral from the lesion site are shown. High magnification insets show select *ctgfa*-Tracer^+^ cells in triple, double, or single channel views. White arrowheads indicate *ctgfa*-Tracer^+^ HuC/D^+^ cells; yellow arrowheads indicate *ctgfa*-Tracer^-^ HuC/D^+^ cells. **(C**,**D)** Quantification of *ctgfa*-Tracer^+^ and HuC/D^+^ colocalization at 28 dpi. *ctgfa*-Tracer Hoechst and HuC/D Hoechst were used to quantify the numbers of *ctgfa*-Tracer^+^ and HuC/D^+^ cells. Percent *ctgfa*-Tracer^+^ HuC/D^+^ cells were normalized to the total number of *ctgfa*-Tracer^+^ cells in C, and to the total number of HuC/D^+^ cells in D. **(E)** Immunostaining for mCherry (green), Mbp (magenta), and Hoechst (grey) at 28 dpi. SC sections from TAM-treated *ctgfa*-Tracer (Cre^+^) animals are shown. Cross sections 150 µm (Proximal) or 450 µm (Distal) rostral from the lesion site are shown. High magnification insets show select *ctgfa*-Tracer^+^ cells in triple, double, or single channel views. White arrowheads indicate *ctgfa*-Tracer^+^ Mbp^+^ staining; yellow arrowheads indicate *ctgfa*-Tracer^+^ Mbp^-^ staining. **(F**,**G)** Quantification of *ctgfa*-Tracer^+^ and Mbp^+^ colocalization at 28 dpi. *ctgfa*-Tracer^+^ and Mbp^+^ fluorescence were quantified. Percent *ctgfa*-Tracer^+^ Mbp^+^ was normalized to total *ctgfa*-Tracer^+^ in F and to total Mbp^+^ fluorescence in G. For all quantifications, SC cross sections 150, 450, and 750 µm rostral to the lesion were analyzed. Dots indicate individual animals and sample sizes are indicated in parentheses. Scale bars, 50 µm.

### Mapping the gene regulatory sequences that direct *ctgfa* expression after SCI

EMT activation localizes to ventral glial progenitors after SCI. The EMT-driving gene regulatory network that includes *ctgfa* is necessary and sufficient to promote glial bridging and functional SC repair (Mokalled et al., 2016; Klatt Shaw et al., 2021). To better understand the regulatory mechanisms that induce localized *ctgfa* expression and EMT activation after SCI, we analyzed the *cis*-regulatory elements that direct injury-induced *ctgfa* transcription in regenerating SC tissues. Using *-5*.*5Kb-ctgfa*:EGFP, we previously showed that EGFP fluorescence recapitulates endogenous mRNA expression in ventral glial progenitors and in bridging glia during SC regeneration (Mokalled et al., 2016). To identify specific regulatory elements within the −5.5 Kb genomic region of *-5*.*5Kb-ctgfa*:EGFP, stable transgenic lines expressing an EGFP cassette downstream of the −4 Kb- or −3 Kb genomic regions were generated (*-4kb-ctgfa*:EGFP and *-3kb-ctgfa*:EGFP, respectively) (Fig. 3A). Multiple independent lines were generated for each transgene to control for possible positional effects that may impact transgene expression. To establish stable reporter lines, transgenesis was first confirmed by performing EGFP PCR, and then by assessing EGFP fluorescence in developing zebrafish embryos (Fig. S3). At 3 days post-fertilization, we observed comparable EGFP expression in *-5*.*5kb-, −4kb-*, and *-3kb-ctgfa*:EGFP. To determine the expression of *ctgfa* reporter lines after SCI, adult reporter animals were subjected to complete SC transections and examined for EGFP expression at 10 dpi (Fig. 3B). *ctgfa*-driven EGFP and Gfap expression were assessed 150 µm (proximal) and 450 µm (distal) rostral to the lesion. Similar to *-5*.*5Kb*-*ctgfa*:EGFP, *-4Kb-ctgfa*:EGFP transgenic lines showed EGFP expression at 10 dpi, while EGFP was not detectable in *-3Kb-ctgfa*:EGFP lines (Fig. 3C). These results suggested that sequences contained within a 971 bp *cis*-regulatory element (referred to hereafter as 1 Kb enhancer element) located between 3 and 4 Kb upstream of the *ctgfa* translational start site are sufficient to direct *ctgfa* expression after SCI.

**Figure 3.**
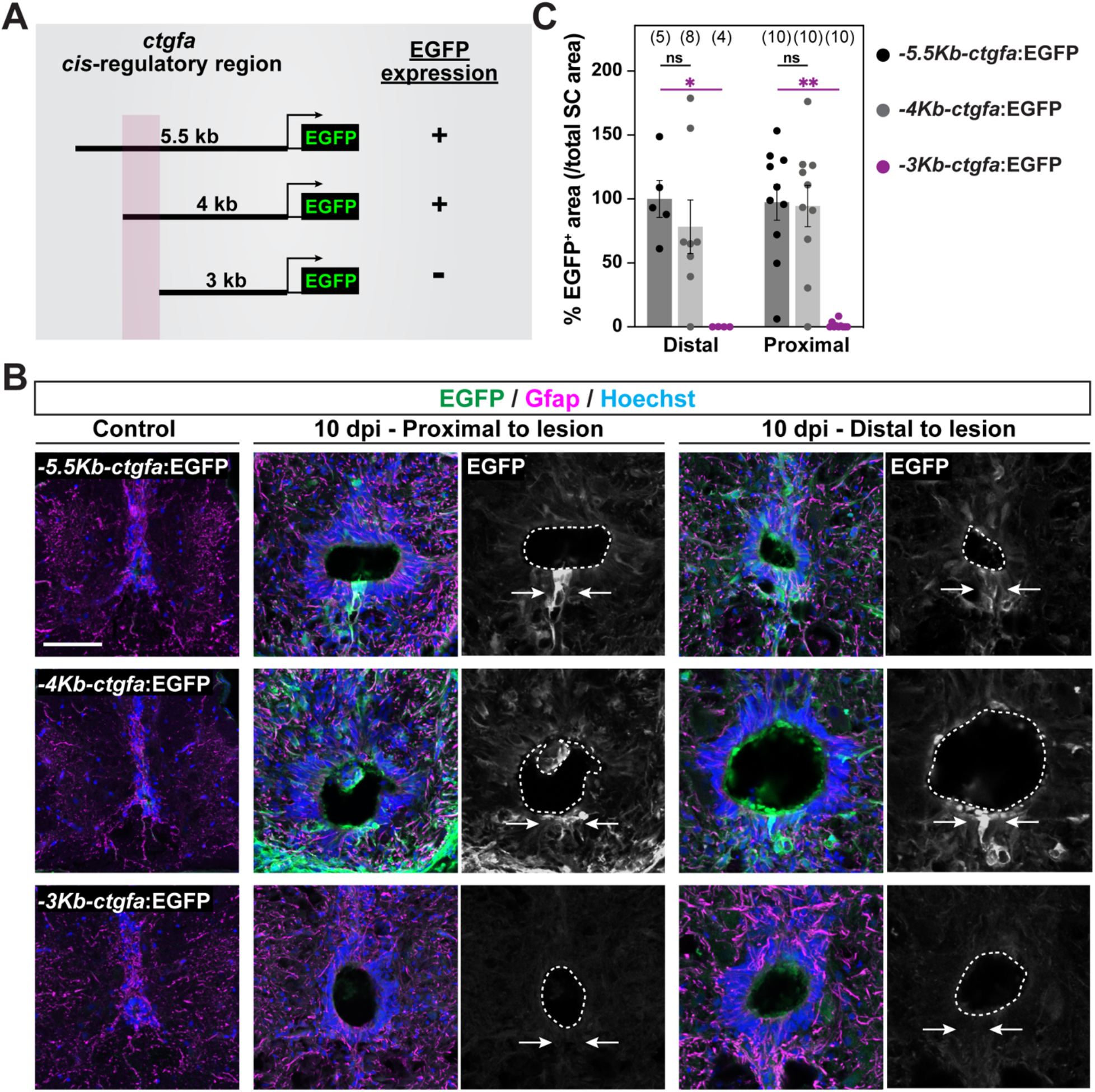
Regulation of *ctgfa* expression during SC regeneration. **(A)** A series of transgene constructs were used to identify the *cis*-regulatory region that drives *ctgfa*-dependent EGFP expression after SCI. A minimum of 3 stable lines were generated for each transgene. **(B)** EGFP and Gfap immunostaining assessed reporter expression at 10 dpi and in uninjured control tissues. SC tissues from *-5*.*5Kb-ctgfa*:EGFP, *-4Kb-ctgfa*:EGFP and *-3Kb-ctgfa*:EGFP transgenic animals are shown. Cross sections are shown at 150 µm (proximal) and 450 µm (distal) rostral to the lesion. Dotted lines delineate central canal edges. Arrows point to EGFP expression in ventral ependymal progenitors. **(C)** Quantification of EGFP fluorescence in *-5*.*5Kb-ctgfa*:EGFP, *-4Kb-ctgfa*:EGFP and *-3Kb-ctgfa*:EGFP transgenic animals. SC cross sections 150 µm (proximal) and 450 µm (distal) rostral to the lesion were quantified. Dots indicate individual animals and sample sizes are indicated in parentheses. \*\**P*<0.01; *P<0.05; ns indicates P>0.5. Scale bars, 50 µm.

### A distal enhancer element directs glial *ctgfa* expression after SCI

To test whether the putative 1 Kb enhancer element is sufficient to promote *ctgfa*-driven EGFP expression after SCI, we generated *1Kb-ctgfa*:EGFP transgenic lines (Fig. 4A). We first tested whether larval EGFP expression in this line recapitulated *-5*.*5Kb*-, *-4Kb*- and *-3Kb*-*ctgfa*:EGFP expression at 3 days post-fertilization, finding that it did (Fig. S3). To minimize the impact of positional effects on transgene expression, 3 independent *1Kb-ctgfa*:EGFP lines were generated and referred to as L1, L2, and L3. SC tissues from adult *1Kb-ctgfa*:EGFP transgenic fish were then transected to examine EGFP expression following injury. At 10 dpi, EGFP was not detectable in uninjured SC tissues, but was comparable to *-5*.*5Kb-* and *-4Kb-ctgfa*:EGFP expression in SC sections 150 µm (proximal) and 450 µm (distal) rostral to the lesion (Fig. 4B,C). This expression pattern was recapitulated in lines L1, L2, and L3. Bioinformatics analysis of the 1 Kb genomic region that directs *ctgfa* expression after injury revealed predicted binding sites for Tead (3 binding sites), JunB (2 binding sites) and Spi1 (6 binding sites) transcription factors (Fig. S4A). Tead factors, which associate with the Yap/Taz co-activators to control gene transcription, are known upstream regulators of *ctgfa* expression in multiple systems. Consistent with their role in promoting cell proliferation and stem cell maintenance (Vassilev et al., 2001; Zhao et al., 2008), *in situ* hybridization showed *ctgfa* expression was downregulated upon transient Yap/Taz knockdown in injured SC tissues (Fig. S4B). *ctgfa* expression was also downregulated in *junbb* CRISPR mutants (Fig. S4C), but not in *spi1a* mutants (Fig. S4D). These results indicated a 1 Kb enhancer element directs injury-induced *ctgfa* expression during SC regeneration as well as larval expression, and suggested *ctgfa* expression after SCI is induced by cooperative regulation of Tead and JunB transcription factors.

**Figure 4.**
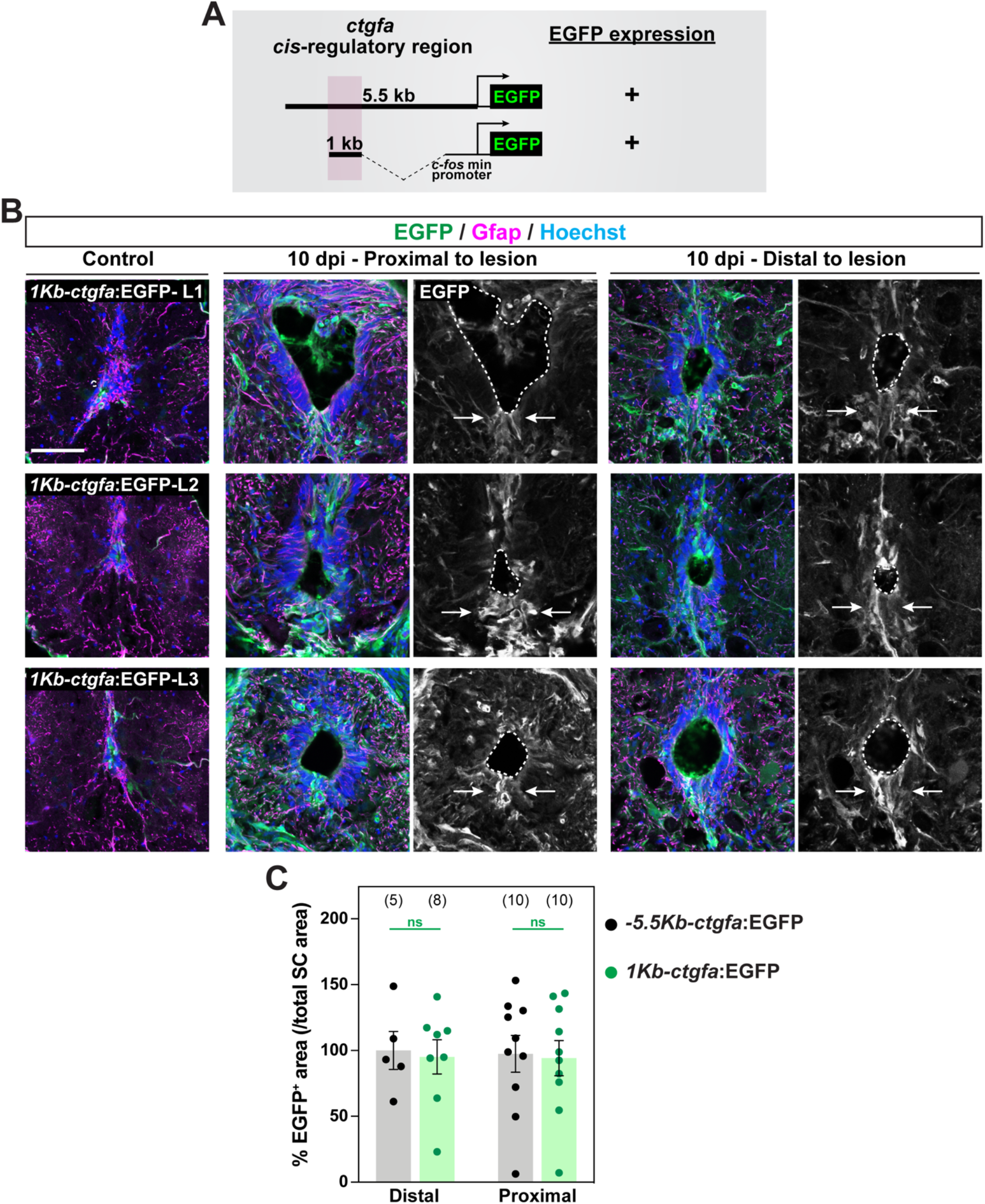
A distal enhancer element directs glial *ctgfa* expression after SCI. **(A)** Sequences within a 1 Kb enhancer element located 3 to 4 Kb were combined with a 96 bp mouse *c-fos* minimal and subcloned upstream of an EGFP-expressing cassette. The clone was co-injected into one-cell stage wild-type embryos, and 3 founders were isolated for propagation. **(B)** EGFP and Gfap immunostaining assessed reporter expression at 10 dpi and in uninjured tissue. SC tissues from three independent lines of *1Kb-ctgfa*:EGFP transgenic animals (L1, L2, and L3) are shown. Cross sections are shown at 150 (proximal) and 450 (distal) µm rostral to the lesion. Dotted lines delineate central canal edges. Arrows point to EGFP expression in ventral ependymal progenitors. **(C)** Quantification of EGFP fluorescence in *-5*.*5Kb-ctgfa*:EGFP and *1Kb-ctgfa*:EGFP-L1 transgenic animals. SC cross sections 150 µm (proximal) and 450 µm (distal) rostral to the lesion were quantified. Dots indicate individual animals and sample sizes are indicated in parentheses. ns indicates P>0.5. Scale bars, 50 µm.

### Ablation of *ctgfa*^+^ cells impairs SC regeneration

Previous studies showed that *ctgfa* mutations impair glial bridging, axon regrowth and functional SC repair in zebrafish (Mokalled et al., 2016). However, since Ctgfa is a secreted matricellular molecule, it remained unclear from these reverse genetic studies whether *ctgfa* expressing glial cells are required for SC regeneration in zebrafish. To address this outstanding question, we employed previously generated *ctgfa*:mCherry-Nitroreductase (*ctgfa*:mCherry-NTR) transgenic animals for cell ablation studies (Klatt Shaw et al., 2021). In this system, *ctgfa*-driven expression of the bacterial NTR enzyme was used to catalyze the reduction of the prodrug Metrodinazole (MTZ) into a cytotoxic product that induces cell death (Curado et al., 2008). To ablate *ctgfa*^+^ progenitors and early bridging glia, we subjected *ctgfa*:mCherry-NTR (Tg^+^) fish and their wild-type (Tg^-^) siblings to complete SC transections, followed by two consecutive treatments with 0.1 mM MTZ for 24 hrs at 4 and 7 dpi (Fig. 5A). Glial bridging, axon regrowth, swim endurance and swim behavior assays were then performed at 14 dpi to determine the impact of *ctgfa*^+^ cell ablation during SC regeneration (Fig. 5A). For glial bridging, Gfap immunostaining was used to determine the areas of glial bridges at the lesion site relative to the areas of intact SC tissues (Fig. 5B,C). Percent bridging at the lesion site averaged 20% in MTZ-treated Tg^-^ controls, and decreased to 6% in MTZ-treated Tg^+^ animals (Fig. 5D). For anterograde axon tracing, Biocytin was applied at the hindbrain level and Biocytin-labeled axons were traced at 600 µm (proximal) and 1500 µm (distal) caudal to the lesion (Fig. 5E,F). In this assay, axon regrowth was attenuated by 68% in proximal SC tissues from MTZ-treated Tg^+^ animals compared to MTZ-treated Tg^-^ controls (Fig. 5G). These results indicated *ctgfa*^+^ cells are required for the cellular regeneration of glial and axonal bridges after SCI.

**Figure 5.**
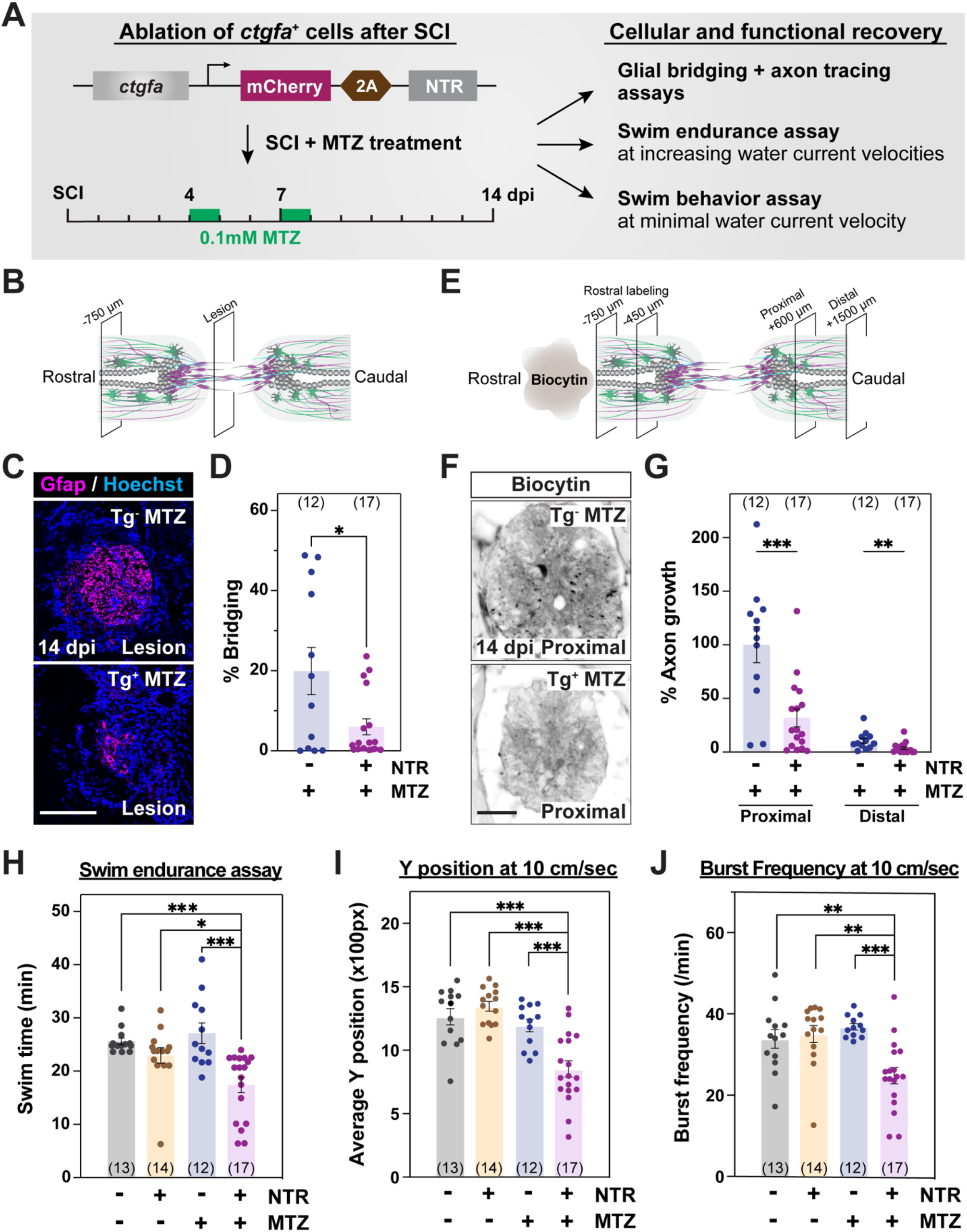
Ablation of *ctgfa*^+^ cells during SC regeneration. **(A)** Experimental timeline to evaluate the requirement of *ctgfa*^+^ cells after SCI. Transgenic *ctgfa:*mCherry-2A-NTR animals and control siblings were subjected to complete SC transection and Metronidazole (MTZ) treatment at 4 and 7 dpi to ablate *ctgfa*^+^ cells. Swim assays were performed and SC tissues were collected at 14 dpi to assess regeneration. **(B-D)** Glial bridging in *ctgfa:*mCherry-2A-NTR animals (Tg^+^) and wild-type siblings (Tg^-^) at 14 dpi. Tg^+^ and Tg^-^ animals were treated with MTZ. Representative immunohistochemistry shows the Gfap^+^ bridge at the lesion site in C. Percent bridging represents the cross-sectional area of the glial bridge at the lesion site relative to the intact SC 750 µm rostral to the lesion in D. **(E-G)** Anterograde axon tracing in *ctgfa:*mCherry-2A-NTR animals (Tg^+^) and wild-type siblings (Tg^-^) at 14 dpi. Biocytin axon tracer was applied rostrally and analyzed at 600 µm (proximal) and 1500 µm (distal) caudal to the lesion (E). Representative images show the extent of Biocytin labeling 600 µm (proximal) caudal to the lesion in F. Axon growth was first normalized to the extent of Biocytin at 450 and 750 µm rostral to the lesion, and then to labeling in Tg^-^ controls at the proximal level in G. **(H)** Swim endurance assays for *ctgfa:*mCherry-2A-NTR animals and wild-type siblings at 14 dpi. This behavioral test assessed the capacity of *ctgfa*-ablated animals to swim in an enclosed swim tunnel under increasing water current velocities. The time at exhaustion for each fish is shown. **(I**,**J)** Swim behavior assays assessed the performance of *ctgfa:*mCherry-2A-NTR animals and wild-type siblings at minimal water current velocity at 14 dpi. Average Y position in the tunnel (I) and burst frequency (J) were quantified at 10 cm/sec water current velocity. For all quantifications, dots represent individual animals and animal numbers are indicated in parentheses. ***P<0.001; **P<0.01; and *P<0.05. Scale bars, 100 µm.

To evaluate the functional impact of ablating *ctgfa*^+^ cells after SCI, we first assessed the swim capacities of ablated animals in an enclosed swim tunnel under increasing water current velocities (Fig. 5H) (Burris et al., 2021). In this swim endurance assay, control animals showed comparable swim functions, averaging between 22 min for vehicle-treated Tg^+^ fish and 27 min for MTZ-treated Tg^-^ fish. However, swim endurance was significantly diminished in MTZ-treated Tg^+^ animals, with a swim time average of 17 min. We then tracked the swim behavior of ablated fish under a constant, low current velocity of 10 cm/sec (Burris et al., 2021). Fish position in the swim tunnel (Y position) and burst frequency were quantified to assess overall swim competence. In these swim behavior assays, MTZ-treated Tg^+^ animals stalled in the back quadrant of the swim tunnel (Fig. 5I), and displayed 32% less frequent bursts under low current velocity (Fig. 5J). Together, these experiments indicated that ablation of *ctgfa*^+^ cells impairs cellular and functional recovery after SCI, and that *ctgfa*^+^ glial cells play an essential pro-regenerative role during SC regeneration in zebrafish.

## DISCUSSION

This study explores the contribution, regulation, and requirements of specialized glial cells during innate spinal cord repair in zebrafish. Our genetic lineage tracing and cell ablation results are consistent with a model in which *ctgfa*-expressing progenitor and glial cells contribute to and are required for glial bridging and functional recovery after SCI.

Following SC transection injuries, the severed SC stumps retract away from the lesion core. Rapid activation and proliferation of ependymal progenitors seal the severed central canal at both ends of the lesion. As regeneration proceeds, the SC stumps are attached by newly formed regenerate tissues, the central canal reconnects, and ependymal progenitors regain their epithelial-like central canal-lining morphology. Our genetic lineage tracing studies showed *ctgfa*^+^ cells give rise to regenerating glia, including bridging glia with minimal contribution to neuronal or oligodendrocyte lineages. These results are consistent with a model in which *ctgfa*^+^ progenitors differentiate intro glia after SCI. Our lineage tracing studies employed Tamoxifen labeling at 4 dpi, a time point at which *ctgfa* is highly expressed but not restricted to ventral ependymal progenitors. Specifically, *ctgfa* expression at 4 dpi marks early bridging glia at the lesion site in addition to additional cell types around injured SC tissues. We thus could not rule out the possibilities that *ctgfa*^+^ glia labeled at 4 dpi proliferate and give rise to more bridging glia, or that *ctgfa*^+^ cells around the lesion transdifferentiate into glia during SC regeneration. We also showed that *ctgfa*-driven lineage tracing labels ventral ependymal progenitors at 28 dpi, suggesting *ctgfa*^+^ progenitors undergo self-renewal and contribute to the newly formed ventral ependyma within regenerate tissues. An alternative possibility is that *ctgfa*^+^ glia may undergo dedifferentiation into ependymal progenitors at later stages of regeneration. We propose that devising future strategies that enable differential targeting of *ctgfa*^+^ progenitors and *ctgfa*^+^ glia are needed to better understand the potency and proliferative capacities of progenitor and glial cells during SC regenerating.

Previous studies implicated Fgf and Ctgf signaling in glial bridge formation in adult zebrafish. The glial bridging functions of Fgf signaling were established using transgenic dominant-negative Fgf receptor manipulations, *sprouty* mutants and Fgf8 injection (Goldshmit et al., 2012). *ctgfa* genetic loss-of-function was shown to impair glial bridging, axon regrowth and functional SC repair (Mokalled et al., 2016). However, since Fgf and Ctgf are secreted extracellular proteins, the glial requirements of these molecules during bridging and of glial bridging during SC regeneration remained unclear. Studies from larval zebrafish suggested axon regrowth proceeds independently of the projection of glial processes across the lesion (Briona et al., 2015; Wehner et al., 2017). Specifically, using a *gfap*-driven Nitroreductase-expressing transgenic line, Wehner et al. showed that axon regrowth was unaffected following Gfap^+^ cell ablation in larval zebrafish (Wehner et al., 2017). We here show that ablation of *ctgfa*^+^ cells was sufficient to impair glial bridging, axon regrowth and functional recovery after SCI, indicating *ctgfa*^+^ cells are required during SC repair. Noting the technical limitations of the NTR-MTZ system at simultaneously achieving complete and specific cell ablation, it is possible that a small proportion of Gfap^+^ cells escaped cell ablation in the larval studies by Wehner *et al*., or that our *ctgfa*-driven studies ablated *ctgfa*^+^ glial progenitors in addition to *ctgfa*^+^ bridging glia. However, regenerating axons often associate with elongated bipolar astrocytes across vertebrates, including zebrafish and mammals (White et al., 2008; Filous et al., 2010; Goldshmit et al., 2012; Zukor et al., 2013; Mokalled et al., 2016). Bridging glia from adult zebrafish retain an elongated morphology (Goldshmit et al., 2012; Mokalled et al., 2016), and possess a mesenchymal signature that correlates with increased plasticity and immature astrocytic cell identity (Klatt Shaw et al., 2021). Similarly in adult mice, a small proportion of elongated astrocytes, termed “astroglial bridges”, correlate with increased axon regrowth under genetic manipulations such as PTEN deletion (Zukor et al., 2013). We thus propose that the requirements for axon regrowth may differ between larval and adult zebrafish, and that future comparative studies will shed light into the molecular similarities and differences between murine elongated astrocytes and zebrafish bridging glia.

Our findings shed light on glial bridging as an effective, natural mechanism of SC regeneration and established glial cell responses are pro-regenerative and indispensable to achieve SC repair. We suggest that further investigation into glial cell fates will springboard translational applications to improve bridging and regeneration in the mammalian CNS.

## ACKNOWLEDGMENTS

We thank V. Cavalli, A. Johnson and L. Solnica-Krezel for discussion, S. Kucenas for sharing the Mbp antibody, and the Washington University Zebrafish Shared Resource for animal care. This research was supported by grants from the NIH to K.D.P. (R21 NS124635 and R21 NS096617) and to M.H.M. (R01 NS113915 and R01 NS123708).

## MATERIALS AND METHODS

### Zebrafish

Adult zebrafish of the Ekkwill, Tubingen, and AB strains were maintained at the Washington University Zebrafish Core Facility. All animal experiments were performed in compliance with institutional animal protocols. Male and female animals between 3 and 9 months of age and of ∼2 cm in length were used. Experimental fish and control siblings of similar size and equal sex distribution were used for all experiments. SC transection surgeries and regeneration analyses were performed in a blinded manner, and 2 to 4 independent experiments were repeated using different clutches of animals for all experiments. Transected animals from control and experimental groups were housed in equal numbers (4-7 fish) in 1.1 liter tanks. The following previously published zebrafish strains were used: Tg(*ctgfa*:mCherry-NTR)^*stl650*^ (Klatt Shaw et al., 2021), Tg(*-5*.*5Kb-ctgfa*:EGFP)^*pd96*^ (Mokalled et al., 2016), *junbb*^*stl672*^ (Klatt Shaw et al., 2021) and *spi1a*^*stl669*^ (Klatt Shaw and Mokalled, 2021). *yap1/taz* crispants were generated as previously described (Klatt Shaw and Mokalled, 2021). Newly constructed strains are described below.

#### *Generation of transgenic* Tg(*ctgfa*:CreER^T2^) *zebrafish*

The following primers were used to amplify a 5.5 kb genomic DNA region upstream of the *ctgfa* translational start site: ClaI forward primer 5’-atcgattttggctactttcagctaagactgg-3’ and ClaI reverse primer 5’-atcgattctttaaagtttgtagcaaaaagaaa-3’. The genomic fragment was cloned into PCR2.1-TOPO vector, then subcloned into ClaI digested PCS2-CreER^T2^ plasmid to generate *ctgfa*:CreER^T2^ clone. The clone was co-injected into one-cell stage wild-type embryos with I-SceI. Multiple founders were isolated and propagated. The full name of this line is Tg(*ctgfa*:CreER^T2^)^*stl652*^. *ctgfa*:CreER^T2^ animals were crossed into *ubiquitin:*loxP-GFP-STOP-loxP-mCherry transgene (Mosimann et al., 2011) to generate *ctgfa*:Tracer animals.

#### *Generation of transgenic* Tg(*-4Kb-ctgfa:*EGFP) *and* Tg(*-3Kb-ctgfa:*EGFP) *zebrafish*

The following forward primers were used to amplify 4 and 3 Kb genomic region upstream of the *ctgfa* translational start site: ClaI 4 Kb forward primer 5’-ccatcgataggcagcaatagcgtcagat-3’ and ClaI 3 Kb forward primer 5’-ccatcgattttgacccctctcagtgaa-3’. A common reverse primer was used to amplify 4 and 3 Kb genomic regions: ClaI reverse primer 5’-ccatcgatttctttaaagtttgtagcaaaaaaga-3’. The genomic fragments were cloned into PCR2.1-TOPO vector, then subcloned into ClaI digested PCS2-EGFP plasmid. Clones were co-injected into one-cell stage wild-type embryos with I-SceI. A minimum of 3 founders were isolated and propagated for each transgene. The full names of these lines are Tg*(−4Kb*-*ctgfa*:EGFP)^*stl656*^ and Tg*(−3Kb*-*ctgfa*:EGFP)^*stl657*^. *A*nimals were analyzed as hemizygotes.

#### *Generation of transgenic* Tg(*1Kb-ctgfa:*EGFP) *zebrafish*

The following primers were used to amplify a 1Kb genomic region upstream of the *ctgfa* translational start site: ClaI forward primer 5’-ccatcgatccacaaggctattgcaacg-3’ and ClaI reverse primer 5’-ccatcgatagtatgcacctattcactgag-3’. The genomic fragments were cloned into PCR2.1-TOPO vector, then subcloned into ClaI digested PCS2-fos-EGFP plasmid. The clone was co-injected into one-cell stage wild-type embryos with I-SceI. Three founders were isolated and propagated. The full names of these lines are Tg*(1Kb*-*ctgfa*:EGFP-L1)^*stl658*^, Tg*(1Kb*-*ctgfa*:EGFP-L2)^*stl659*^, and Tg*(1Kb*-*ctgfa*:EGFP-L3)^*stl660*^. Animals were analyzed as hemizygotes.

### Spinal cord transection and treatment

Zebrafish were anaesthetized using MS-222. Fine scissors were used to make a small incision that transects the spinal cord 4 mm caudal to the brainstem region. Complete transection was visually confirmed at the time of surgery. Injured animals were also assessed at 2 or 3 dpi to confirm loss of swim capacity post-surgery.

### Histology

Sixteen µm cross cryosections of paraformaldehyde-fixed SC tissues were used. Tissue sections were imaged using a Zeiss AxioVision compound microscope or a Zeiss Axioscan.Z1 slide scanner for *in situ* hybridization, and a Zeiss LSM 800 confocal microscope or a Zeiss Axioscan.Z1 slide scanner for immunofluorescence.

For *in situ* hybridization, a previously cloned *ctgfa* probe was used (Mokalled et al., 2016). Linearized vectors were used to generate the digoxygenin labeled cRNA probes. *in situ* hybridization assays were performed as previously described (Mokalled et al., 2016).

Primary antibodies used in this study were rabbit anti-dsRed (Clontech, 632496, 1:200), mouse anti-GFAP (ZIRC, ZRF1, 1:500), mouse anti-HuC/D (Invitrogen, A21271, 1:1000), mouse anti-Hb9 (Developmental Studies Hybridoma Bank, AB2145209, 1:50), rabbit anti-Mbp (a generous gift from S. Kucenas lab, 1:500), chicken anti-GFP (Aves Labs, GFP-1020, 1:1000). Secondary antibodies (Invitrogen, 1:200) used in this study were Alexa Fluor 488, Alexa Fluor 594, and Alexa 647 goat anti-rabbit or anti-mouse antibodies.

### Quantification

#### Colocalization analysis

Colocalization of *ctgfa*-Tracer with Gfap and of *ctgfa*-Tracer with Mbp were performed using JACoP plugin in Fiji. Orthogonal projections of individual image stacks were generated using Zen software. The polygon selection tool was used to outline the spinal cord perimeters to define regions of interests (ROIs). Thresholds were user-defined for the *ctgfa*-Tracer channel and Gfap/Mbp channel. M1 and M2 coefficients were calculated. M1 is defined as the ratio of the summed intensities of pixels from the *ctgfa*-Tracer channel for which the intensity in the Gfap/Mbp channel is above zero to the total intensity in the Gfap/Mbp channel. M2 is defined as the ratio of the summed intensities of pixels from the *ctgfa*-Tracer channel for which the intensity in the Gfap/Mbp channel is above zero to the total intensity in the *ctgfa*-Tracer channel. M1 and M2 coefficients were multiplied by 100 to obtain percent expression. Two-way ANOVA and multiple comparisons were performed using the Prism software to determine statistical significance of swim times between groups.

#### Cell counting

Cell counting of *ctgfa*-Tracer with HuC/D and of *ctgfa*-Tracer with Hb9 were performed using a customized Fiji script (adapting ITCN: Image based Tool for counting nuclei-https://imagej.nih.gov/ij/plugins/itcn.html). Orthogonal projections of individual image stacks were generated using Zen software. A Customized Fiji script incorporated user-defined inputs to define channels (including Hoechst) and to outline SC perimeters. To quantify nuclei, the following parameters were set in ITCN counter: width, 15; minimal distance, 7.5; threshold, 0.4. Once nuclei were identified, user defined thresholds of individual cell markers were used to mask the image and identify nuclei located inside the masked regions. X/Y coordinates were extracted for each nucleus for cell counting. Raw counts and X/Y coordinates from Fiji were processed using a customized R script. Two markers are considered overlapping if they share nuclei with same X/Y coordinates.

#### Quantification of ctgfa:EGFP expression

Images were thresholded blind to condition, and the areas of the thresholded signal were measured. To calculate percent EGFP^+^ area, the thresholded areas were normalized to total SC area for each section. Unpaired t-tests with Welch’s correction were performed using Prism software to determine statistical significance between groups.

#### Quantification of ctgfa in situ hybridization

Images were converted into a 16-bit tiff and a horizontal line was drawn at the center of the central canal to separate the dorsal and ventral sections of the SC. Images were thresholded blind to condition, and the areas of the thresholded signal were measured in ventral SC tissues. To calculate percent *ctgfa*^+^ area, the thresholded areas were normalized to total SC area for each section. Unpaired t-tests with Welch’s correction were performed using Prism software to determine statistical significance between groups.

### Glial bridging

GFAP immunohistochemistry was performed on serial transverse sections. The cross-sectional area of the glial bridge (at the lesion site) and the area of the intact SC (750 µm rostral to the lesion) were measured using ImageJ software. Bridging was calculated as a ratio of these measurements. Unpaired t-tests with Welch’s correction were performed using Prism software to determine statistical significance between groups.

### Axon tracing

Anterograde axon tracing was performed on adult fish at 28 dpi. Fish were anaesthetized using MS-222 and fine scissors were used to transect the cord 4 mm rostral to the lesion site. Biocytin-soaked Gelfoam Gelatin Sponge was applied at the new injury site (Gelfoam, Pfizer, cat# 09-0315-08; Biocytin, saturated solution, Sigma, cat# B4261). Fish were euthanized 6 hours post-treatment and Biocytin was histologically detected using Alexa Fluor 594-conjugated Streptavidin (Molecular Probes, cat# S-11227). Biocytin-labeled axons were quantified using the “threshold” and “particle analysis” tools in the Fiji software. Four sections per fish at 600 µm (proximal) and 1500 µm (distal) caudal to the lesion core, and 2 sections 450 and 750 µm rostral to the lesion, were analyzed. Axon growth was normalized to the efficiency of Biocytin labeling rostral to the lesion for each fish. Percent axon growth was then normalized to the rostral level of the control group. Unpaired t-tests with Welch’s correction were performed using Prism software to determine statistical significance between groups.

### Swim endurance assays

Zebrafish were exercised in groups of 8-12 in a 5 L swim tunnel device (Loligo, cat# SW100605L, 120V/60Hz). After 10 minutes of acclimation inside the enclosed tunnel, water current velocity was increased every two minutes and fish swam against the current until they reached exhaustion. Exhausted animals were removed from the chamber without disturbing the remaining fish. Swim times at exhaustion were recorded for each fish. Results were expressed as means ± SEM. An unpaired two-tailed Student’s t-test with Welch correction was performed using the Prism software to determine statistical significance of swim times between groups.

### Swim behavior assays

Zebrafish were divided into groups of 5 in a 5 L swim tunnel device (Loligo, cat# SW100605L, 120V/60Hz). Each group was allowed to swim for a total of 15 min under zero to low current velocities (5 min at 0 cm/s, 5 min at 10 cm/s, and 5 min at 15 cm/s). The entire swim behavior was recorded using a high-speed camera (iDS, USB 3.0 color video camera) with following settings: aspect ratio, 1:4; pixel clock, 344; frame rate, 70 frames/s; exposure time: 0.29; aperture, 1.4 to 2; maximum frames; 63,000. Movies were converted to 20 frames/s and analyzed using a customized Fiji macro. For each frame, animals/objects > 1500 px^2^ were identified, and the XY coordinates were derived for each animal/object. Frame were independently, and animal/object tracking was completed using a customized R Studio script. The script aligned coordinates and calculated swim metrics considering three separate frame windows (Frames 0-6000 at 0 cm/s; frames 6001-12000 at 10 cm/s, and frames 12001-18001 at 20 cm/s). One-way ANOVA and multiple comparisons were performed using the Prism software to determine statistical significance of swim times between groups.

