## Supplemental Figures for "Progenitor derived glia are required for spinal cord regeneration in zebrafish"

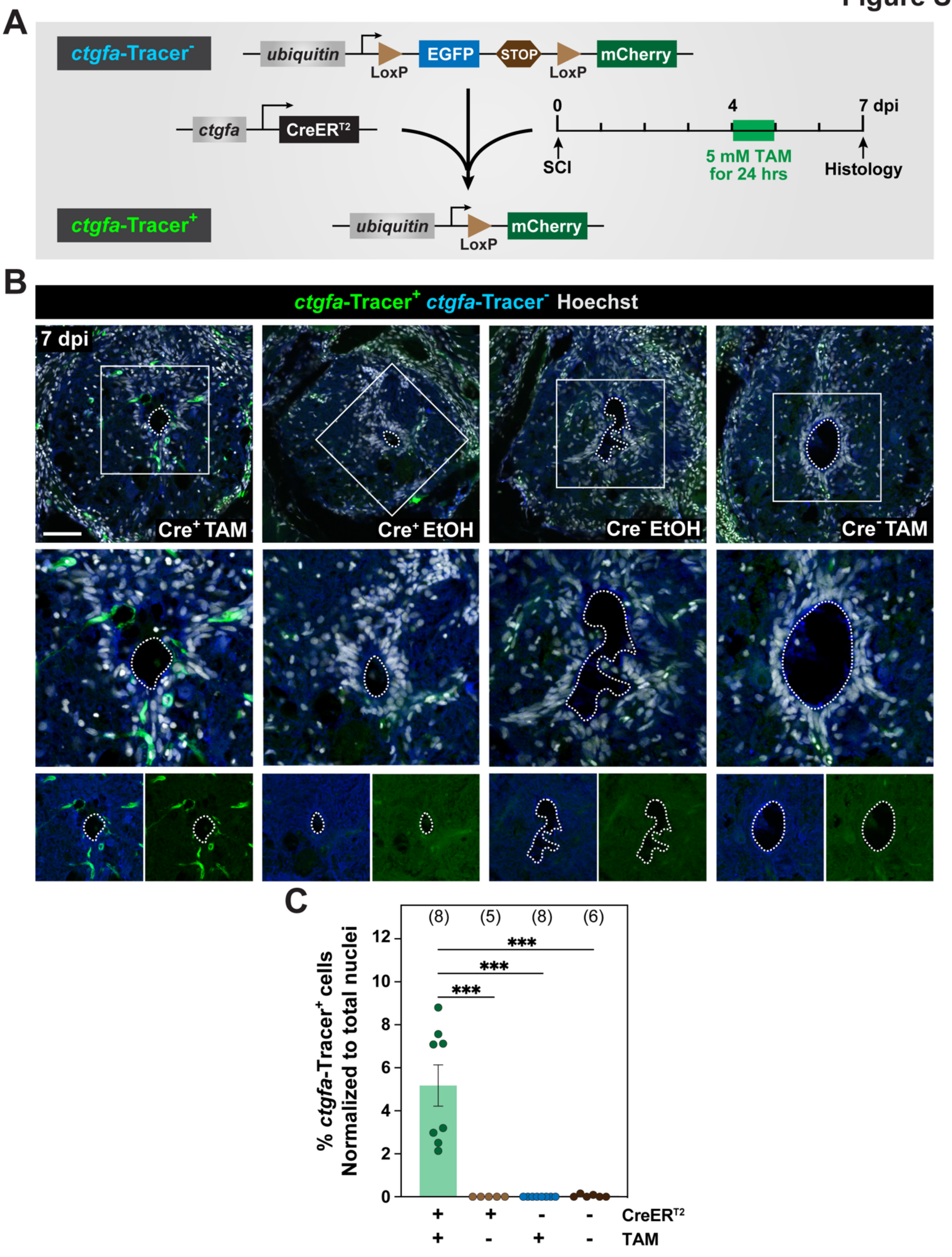
A
ctgfa-Tracer-
ubiquitin
LoxP
EGFP
STOP
LoxP
mCherry
ctgfa
CreER$^{T2}$
0
4
7 dpi
SCI
5 mM TAM for 24 hrs
Histology
ctgfa-Tracer+
ubiquitin
LoxP
mCherry
B
ctgfa-Tracer+ ctgfa-Tracer- Hoechst
7 dpi
Cre+ TAM
Cre+ EtOH
Cre- EtOH
Cre- TAM
C
% ctgfa-Tracer+ cells Normalized to total nuclei
(8)
(5)
(8)
(6)
***
***
***
0
2
4
6
8
10
12
CreER$^{T2}$
TAM
+ + - -
+ - + -

**Figure S1. Generation and characterization of *-5.5Kb-ctgfa*:CreER$^{T2}$ transgenic line. (A)** schematic representation of the generation of *ctgfa*-Tracer zebrafish. A *-5.5Kb-ctgfa:*CreER$^{T2}$ line was generated in combination *ubiquitin:*loxP-GFP-STOP-loxP-mCherry transgene. SC transections were performed on *ctgfa*-Tracer animals and control CreER$^{T2-}$ siblings and treated with either 5 mM Tamoxifen (TAM) or ethanol vehicle (EtOH) at 4 dpi. SC tissues were collected 2 days following the end of Tamoxifen or vehicle treatment, which corresponds to 7 dpi, for histological examination to assess the extent of recombination. **(B)** Immunostaining for mCherry (*ctgfa*-Tracer$^{+}$ in green), GFP (ctgfa-Tracer$^{-}$ in blue), and Hoechst (grey) at 7 dpi. SC sections from *ctgfa*-Tracer (CreER$^{T2+}$) and control siblings (CreER$^{T2-}$) are shown. Treatment conditions include Tamoxifen (TAM) or ethanol vehicle (EtOH). Cross sections 450μm from the lesion site are shown. Dotted line demarcates the central canal. Single channel insets show *ctgfa*-Tracer$^{+}$ and *ctgfa*-Tracer$^{-}$ cells at a higher magnification. **(C)** Quantification of the numbers of *ctgfa*-Tracer$^{+}$ cells at 7 dpi. Percent *ctgfa*-Tracer$^{+}$ cells were normalized to the total number of nuclei for each section. SC cross sections 450 μm rostral to the lesion were analyzed. Dots indicate individual animals and sample sizes are indicated in parentheses. ***P<0.001. Scale bars, 50 μm.

Figure S2

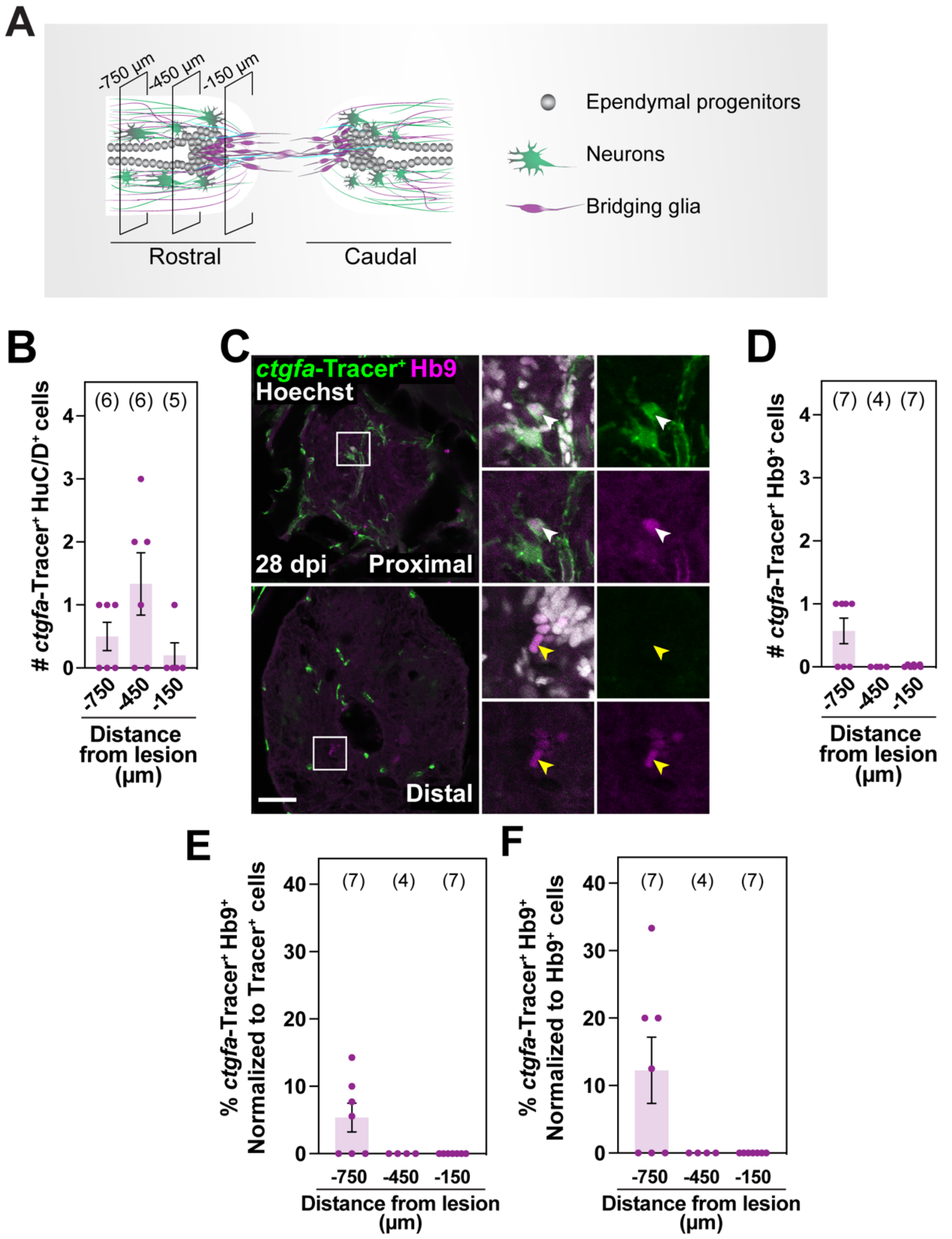
A
-750 µm
-450 µm
-150 µm
Rostral
Caudal
Ependymal progenitors
Neurons
Bridging glia
B
### ctgfa-Tracer+ HuC/D+ cells
(6) (6) (5)
-750 -450 -150
Distance from lesion (µm)
C
ctgfa-Tracer+ Hb9
Hoechst
28 dpi
Proximal
Distal
D
### ctgfa-Tracer+ Hb9+ cells
(7) (4) (7)
-750 -450 -150
Distance from lesion (µm)
E
% ctgfa-Tracer+ Hb9+ Normalized to Tracer+ cells
(7) (4) (7)
-750 -450 -150
Distance from lesion (µm)
F
% ctgfa-Tracer+ Hb9+ Normalized to Hb9+ cells
(7) (4) (7)
-750 -450 -150
Distance from lesion (µm)

**Figure S2. Contribution of *ctgfa* expressing cells to regenerating motor neurons during SC regeneration. (A)** Schematic of injured SC tissues. For lineage tracing studies, tissue sections 150, 450 and 750 μm rostral to the lesion were analyzed. **(B)** Quantification of *ctgfa*-Tracer$^{+}$ and HuC/D$^{+}$ colocalization at 28 dpi. *ctgfa*-Tracer Hoechst and HuC/D Hoechst colocalization were used to quantify the numbers of *ctgfa*-Tracer$^{+}$ and HuC/D$^{+}$ cells. Absolute numbers of *ctgfa*-Tracer$^{+}$ HuC/D$^{+}$ cells per section are shown. **(C)** Immunostaining for mCherry (green), Hb9 (magenta), and Hoechst (grey) at 28 dpi. SC sections from TAM-treated *ctgfa*-Tracer (CreER$^{T2+}$) animals are shown. Cross sections 150 μm (Proximal) or 450 μm (Distal) rostral from the lesion site are shown. High magnification insets show select *ctgfa*-Tracer$^{+}$ cells in triple, double, or single channel views. White arrowheads indicate *ctgfa*-Tracer$^{+}$ Hb9$^{+}$ cells; yellow arrowheads indicate *ctgfa*-Tracer$^{-}$ Hb9$^{+}$ cells. **(D-F)** Quantification of *ctgfa*-Tracer$^{+}$ and Hb9$^{+}$ colocalization at 28 dpi. *ctgfa*-Tracer Hoechst and Hb9 Hoechst colocalization were used to quantify the numbers of *ctgfa*-Tracer$^{+}$ and Hb9$^{+}$ cells. Absolute numbers of *ctgfa*-Tracer$^{+}$ Hb9$^{+}$ cells are shown in D. Percent *ctgfa*-Tracer$^{+}$ Hb9$^{+}$ cells were normalized to the total number of *ctgfa*-Tracer$^{+}$ cells in E, and to the total number of Hb9$^{+}$ cells in F. For all quantifications, dots represent individual animals and animal numbers are indicated in parentheses. Scale bars, 50 μm.

#### Figure S3

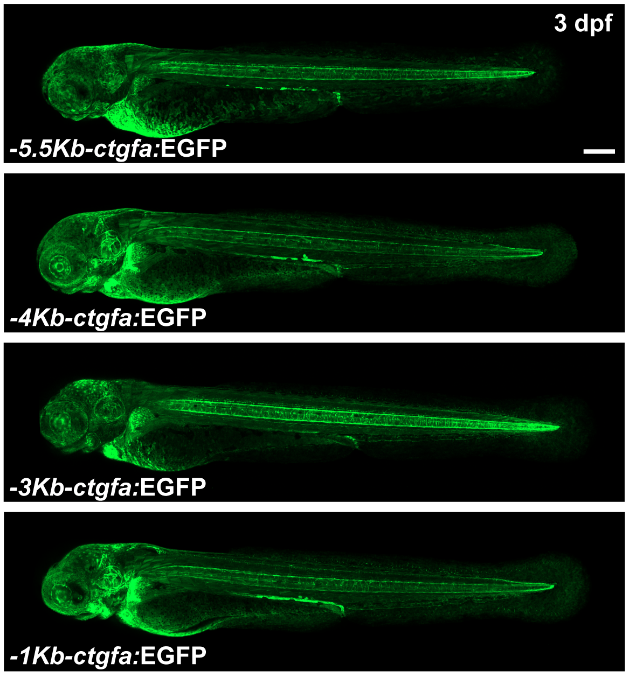


**Figure S3. A 1Kb enhancer element directs *ctgfa* expression during development. (A)** Generation of a series of *ctgfa* reporter lines to identify the *cis*-regulatory region that drives *ctgfa*-dependent EGFP expression after SCI. GFP expression in stable lines from *-5.5Kb-ctgfa*:EGFP, *-4Kb-ctgfa*:EGFP, *-3Kb-ctgfa*:EGFP, and *1Kb-ctgfa*:EGFP(L1) transgenic animals are shown at 3 days post-fertilization (dpf). Scale bars, 200 μm.

**Figure S4**

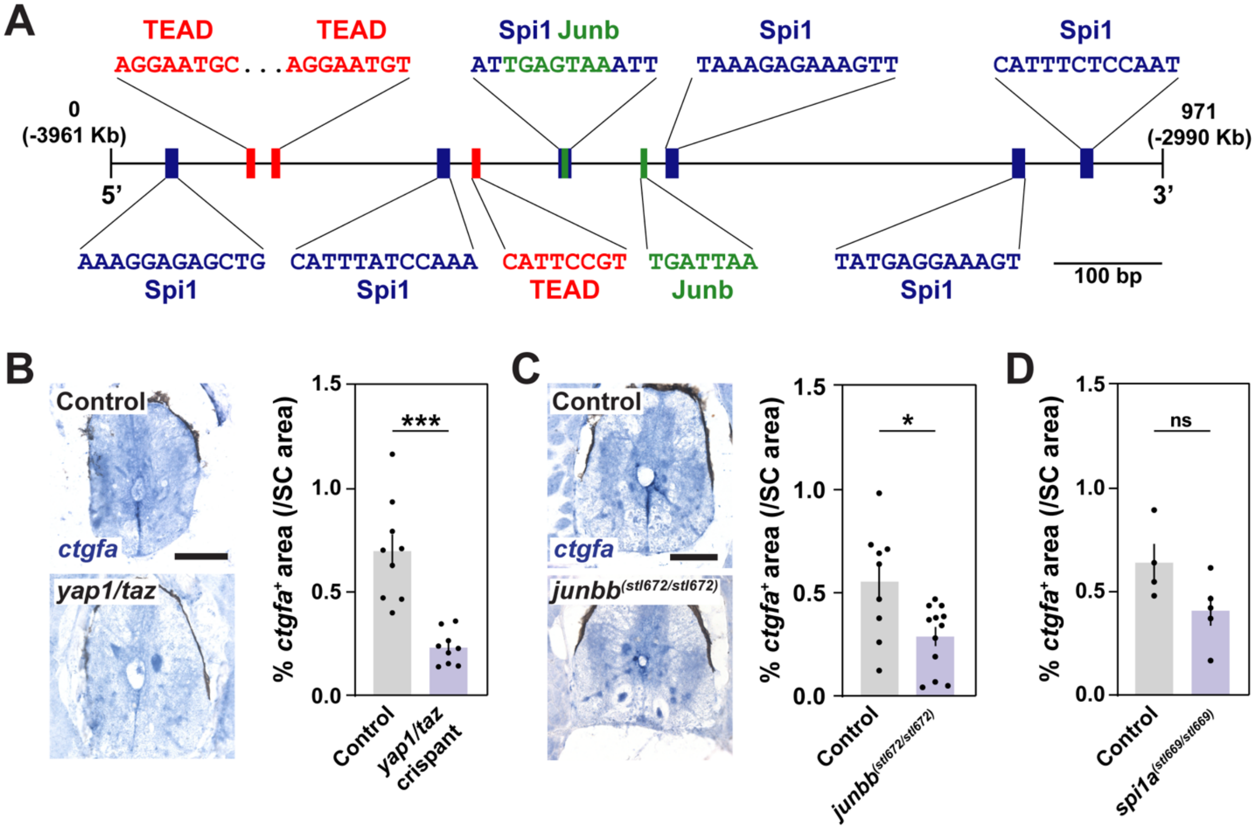


**Figure S4. Regulation of ctgfa expression after SCI. (A)** Transcription factor binding sites within the *1Kb-ctgfa* distal enhancer element. Binding sites for TEAD, Spi1, and Junb transcription factors are indicated. **(B-D)** *in situ* hybridization for *ctgfa* in *yap1/taz* crispants, *junbb* and *spi1a* mutants at 28 dpi. SC cross sections are shown. Uninjected siblings were used as controls for *yap1/taz* crispants. Wild-type siblings were used as controls for *junbb* and *spi1a* mutants. *ctgfa*$^{+}$ areas were quantified 450 μm rostral to the lesion. Percent *ctgfa*$^{+}$ areas were normalized to total SC area for each section. For all quantifications, dots represent individual animals and animal numbers are indicated in parentheses. Scale bars, 100 μm.
